# Chronic alcohol induces subcircuit-specific striatonigral plasticity shifting action control to the sensorimotor striatum

**DOI:** 10.1101/2023.11.15.567225

**Authors:** Giacomo Sitzia, Sebastiano Bariselli, Alexa Gracias, David M. Lovinger

**Affiliations:** Laboratory for Integrative Neuroscience, National Institute on Alcohol Abuse and Alcoholism, National Institutes of Health, Rockville, MD, USA; Department of Neuroscience, Faculty of Health and Medical Sciences, University of Copenhagen, Copenhagen, Denmark; IRCCS Humanitas Research Hospital, Via Manzoni 56, 20089 Rozzano, Milan, Italy

## Abstract

While cortico-striatal circuit deficits contribute to Alcohol Use Disorder, the impact of alcohol on synaptic function in the basal ganglia output, the substantia nigra pars reticulata (SNr), remains unclear. Here, we found that the inputs from the dorsomedial (DMS) and dorsolateral striatum (DLS) differ in their presynaptic properties and target molecularly distinct subpopulations of SNr neurons. We also discovered that indirect pathway subthalamic (STN) inputs to the medial and lateral SNr have different presynaptic properties and that STN inputs are stronger in the lateral SNr. Chronic alcohol exposure (CIE) potentiated DLS inputs but did not affect the strength and presynaptic release properties of DMS and subthalamic inputs to SNr neurons. Chemogenetic inhibition of DLS direct pathway projection neurons impaired action performance in an operant conditioning task in CIE mice but not control mice. Overall, our work identifies a synaptic mechanism whereby chronic alcohol induces a gain of function for action control in direct pathway neurons in the dorsolateral striatum.

**Teaser:** Chronic alcohol selectively potentiates DLS synaptic inputs to the SNr, enhancing their role in action control.

## Introduction

Alcohol is one of the most abused drugs worldwide. Alcohol Use Disorder (AUD) is a disease characterized by loss of control over alcohol intake^1,2^, with devastating consequences for the affected people and society. Alcohol can affect brain function through acute, direct interactions with multiple substrates including, but not limited to, neurotransmitter receptors^3,4^. Alcohol also produces long-term changes in neural circuits^3,5^. Therefore, disentangling the complexity of brain circuits and their susceptibility to alcohol effects is necessary to refine therapeutic strategies targeting behavioral dysfunctions in AUD.

Brain regions of the basal ganglia (BG) coordinate motor and non-motor behaviors based on the present context and past experiences^6–9^, and their dysfunction has been implicated in the development of pathological habits and altered control of goal-directed behavior in AUD^10^. BG circuits control diverse functional outputs through the GABAergic projection neurons of the substantia nigra pars reticulata (SNr) and the globus pallidus internal segment (GPi), which target brainstem, midbrain and thalamic nuclei^11–13^. The BG control the SNr via striatal GABAergic direct pathway inputs and subthalamic nucleus (STN) glutamatergic indirect pathway inputs, that according to the classical view can suppress or promote behavioral outputs influenced by the SNr, respectively^8,9,14^. The main neuronal population of the dorsal striatum (DS) are the spiny projection neurons (SPNs), divided into direct pathway SPNs (dSPNs) targeting the SNr/GPi and indirect pathway SPNs (iSPNs) targeting the globus pallidus external segment (GPe). The GPe in turn projects to the STN and the SNr. The direct and indirect pathway are a circuit backbone shared across anatomically parallel BG circuits with distinct functional roles^15–17^. The dorsomedial (DMS) and dorsolateral (DLS) subregions of the striatum receive associative and sensorimotor cortical inputs^18,19^ and have been implicated in goal-directed (DMS) and habitual (DLS) action control^6^, respectively. Striatal projections to the SNr are topographically organized, hence spatially distinct subpopulations in the medial (^M^SNr) and ventrolateral (^L^SNr) SNr receive inputs from the DMS and DLS^12,20^. Understanding how alcohol affects BG circuits requires a refined knowledge of the organization of synaptic inputs to subpopulations of SNr neurons.

The intrinsic physiological properties of SNr neurons vary along the mediolateral axis of the nucleus^13^. Molecularly identified subpopulations of SNr neurons include a majority of parvalbumin (PV^+^) expressing neurons, enriched in the ventrolateral aspects of the nucleus, as well as other subpopulations that have been characterized based on the expression of the vesicular GABA transporter (VGAT), calretinin (CR) and Glutamate Decarboxylase 2 enzyme (GAD2)^13,21–23^. The physiological properties and projection patterns of PV^+^ SNr neurons distinguish them from other SNr neuron subpopulations^21,24^. The molecular and functional heterogeneity of SNr neurons is also tuned to the diversity of neuronal targets innervated by the SNr^13^. However, whether synaptic inputs to distinct SNr neuron subpopulations are organized to reflect their functional heterogeneity remains unclear.

The synaptic effects of acute and chronic alcohol exposure on the BG have been investigated in animal studies mainly focused on striatal circuits^25,26^. These experiments showed that acute alcohol effects and maladaptive circuit adaptations following chronic alcohol exposure vary according to the striatal cell type, synaptic input, and subregion examined. In a series of seminal studies, Corbit & Janak demonstrated that the control of alcohol self-administration in rats shifts from goal-directed to habitual upon repeated (>2 weeks) exposure due to an increased engagement of the DLS over the DMS^27,28^. Within the DLS, acute exposure of striatal slices to alcohol decreases GABAergic tone on SPNs, inhibits SPN-SPN synapses and fast spiking interneuron-SPN synapses^29^. This suggests that acute alcohol exposure initiates a disinhibition of DLS circuits. Within the DMS, altered synaptic plasticity of the medial prefrontal cortex (mPFC)-DMS inputs contributes to perseveration in alcohol-seeking behaviors^30,31^, whereas weakened orbitofrontal cortex (OFC)-DMS inputs impairs goal-directed action control following chronic alcohol exposure^32,33^. These studies indicate that chronic alcohol exposure induces synaptic maladaptations in specific striatal subcircuits to cause distinct behavioral alterations. However, the susceptibility to alcohol of BG output structures innervated by the striatum remains poorly understood. This limits our understanding of the behavioral and circuit adaptations induced by chronic alcohol exposure.

Here, we asked whether synaptic plasticity in SNr contributes to altered action control following chronic alcohol exposure in mice. First, we used optogenetics-assisted circuit mapping and found that DMS-SNr and DLS-SNr inputs differ in their presynaptic properties and target distinct SNr subpopulations. Moreover, we found that indirect pathway inputs from the STN are biased to the lateral SNr. We then demonstrated that chronic alcohol exposure potentiates DLS-SNr inputs without changing the strength and presynaptic properties of STN-SNr and DMS-SNr inputs. Finally, we found that chemogenetic inhibition of DLS-SNr inputs impaired action execution in mice exposed to chronic alcohol but not in control mice. Overall, we identified a synaptic basis for a shift in action control to the sensorimotor BG circuit following chronic alcohol exposure.

## Results

### 1. Divergent input-output organization of the DMS-SNr and DLS-SNr projections

We virally expressed the Chronos opsin in the DMS or DLS using injections of an adeno-associated virus (AAV) (pAAV5-Syn-Chronos-GFP) and performed *ex vivo* whole-cell patch-clamp recordings coupled with local field optogenetic stimulation in SNr slices (Fig. 1A). As expected, DMS-GFP^+^ terminals were detected in the medial portion of the SNr (^M^SNr). In contrast, DLS-GFP^+^ terminals were detected in the ventrolateral portion of the SNr (^L^SNr)^12,16,20^ (Fig. 1A). Therefore, we focused our recordings on the ^M^SNr to characterize DMS-SNr synapses and on the ^L^SNr to characterize DLS-SNr synapses. The amplitude of oIPSCs at DMS-^M^SNr (2.5 ± 0.6 nA) and DLS-^L^SNr (1.6 ± 0.4 nA) synapses was not statistically different (Fig. supp. 1A), while the oIPSC latency to peak was statistically different but consistent in both synapses with monosynaptic inputs (DMS-^M^SNr 3.8 ± 0.1 ms; DLS-^L^SNr 5.1 ± 0.2 ms) (Fig. supp. 1B). We compared the presynaptic properties of DMS-^M^SNr and DLS-^L^SNr synapses by calculating the paired-pulse ratio (PPR) using stimuli delivered with a 50 ms interpulse interval. We found that PPRs at DMS-SNr synapses were characterized by short-term depression (PPR: 0.87 ± 0.05) and PPRs at DLS-SNr synapses were characterized by short-term facilitation (PPR: 1.29 ± 0.12), and that overall PPRs were significantly different between the two synapses (Fig. 1B). This difference was further evidenced during a train of 10 stimuli delivered at 20 Hz (Fig. supp. 1C). These results indicate that DMS-^M^SNr and DLS-^L^SNr synapses differ in their presynaptic properties.

**Figure 1:**
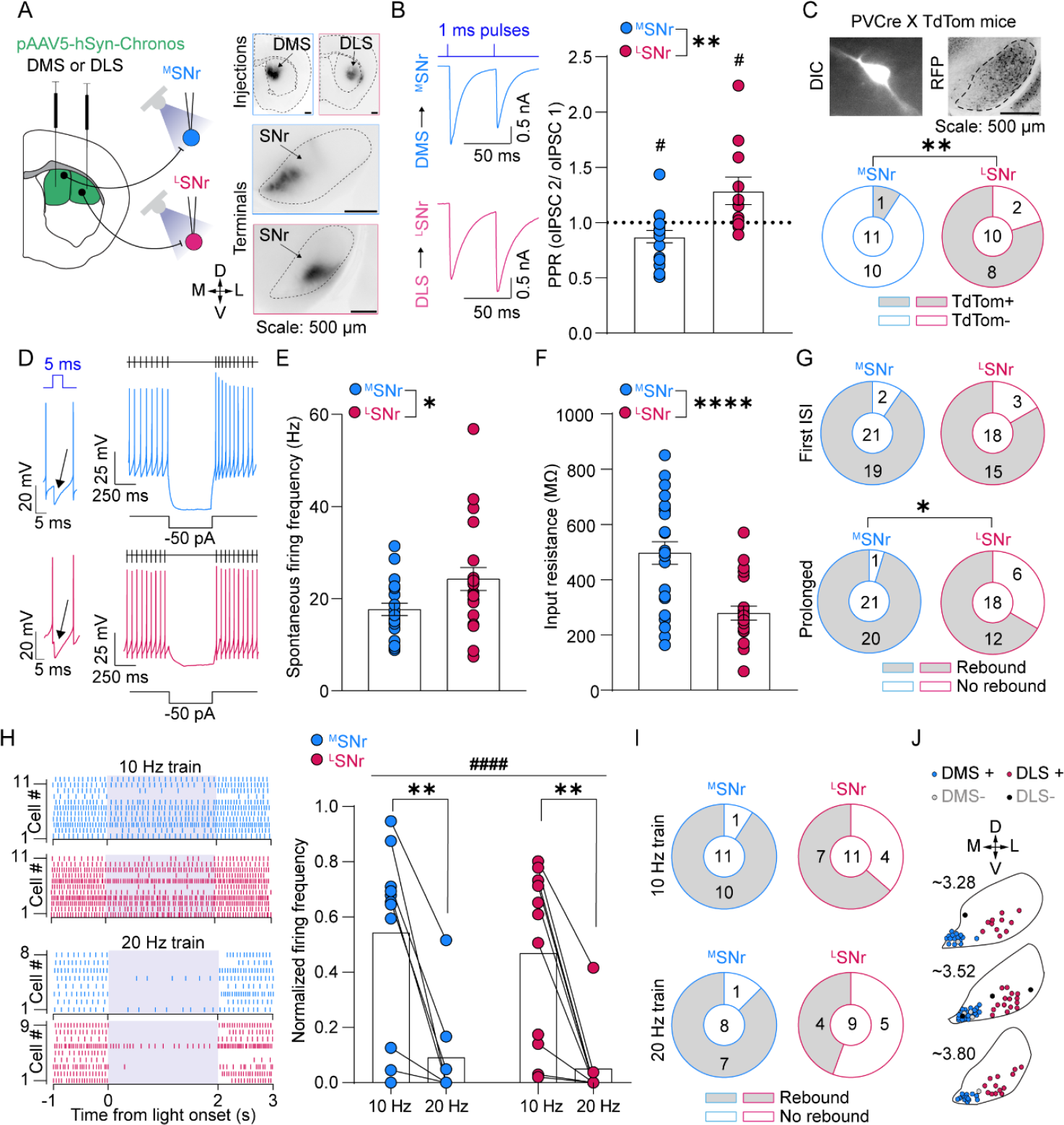
Input-output organization of DMS-SNr and DLS-SNr projections. **A**: Left: Viral injection schematic; Right: validation of Chronos-GFP expression in DMS/ DLS (injection sites, top) and SNr (terminals, bottom). **B**: Paired pulse ratio at DMS-^M^SNr (N = 17 cells, 10 slices, 6 mice) and DLS-^L^SNr synapses (N = 11 cells, 10 slices, 5 mice) (one sample t-test hypothetical value 1, DMS-^M^SNr p = 0,04 and DLS-^L^SNr p = 0.04; Mann-whitney test, two tailed, p = 0.002). **C**: Quantification of PV expressing (TdTom^+^) and not expressing (TdTom^-^) neurons among DMS-targeted (N = 11 cells, 7 slices, 4 mice) and DLS-targeted SNr cells (N = 10 cells, 9 slices, 4 mice) (Fisher’s exact test, p = 0.002). **D**: Example traces and protocols for electrophysiological characterization of DMS-targeted and DLS-targeted SNr neurons. **E**: Spontaneous firing frequency of DMS-targeted (N = 21 cells, 17 slices, 7 mice) and DLS-targeted SNr neurons (N = 21 cells, 14 slices, 6 mice) (Mann-Whitney test, two-tailed, p = 0.04). **F**: Input resistance of DMS-targeted (N = 24 cells, 17 slices, 6 mice) and DLS-targeted (N = 22 cells, 15 slices, 7 mice) SNr neurons (Unpaired t-test, two-tailed, p < 0.0001). **G**: Proportion of ^M^SNr and ^L^SNr neurons displaying fast or prolonged (fisher’s exact test, p = 0.035) rebound responses. **H:** Effects of DMS inputs (N = 11 cells, 10 Slices, 4 mice) and DLS inputs (N = 11 cells, 9 slices, 3 mice) on SNr neuron firing (mixed effects analysis, frequency main effect F_1, 20_ = 25.74, p < 0.0001, Location main effect F_1, 15_ = 0.46; Šídák’s multiple comparisons test, DMS-^M^SNr p = 0.0016, DLS-^L^SNr p = 0.0024). **I**: Proportion of neurons displaying rebound firing following DMS-^M^SNr and DLS-^L^SNr input stimulation at 10 Hz or 20 Hz. **J**: Location of the DMS-targeted and DLS-targeted SNr neurons presented in Fig. 1.

In a subset of these experiments, we used PVCre-TdTom mice to label parvalbumin-expressing (PV^+^) SNr neurons, previously indicated as the major GABAergic subpopulation of the SNr^13,24^. We found a significantly higher proportion of TdTom^-^ neurons (PV^-^) among DMS-connected neurons (1 TdTom^+^, 10 TdTom^-^) compared to a majority of TdTom^+^ neurons (PV^+^) among DLS-connected neurons (8 TdTom^+^, 2 TdTom^-^) (Fig. 1C). This first series of experiments indicated that the input-output organization of DMS-^M^SNr and DLS-^L^SNr projections is divergent, therefore we further investigated their functional properties.

We used the same Chronos-assisted DMS-^M^SNr and DLS-^L^SNr circuit mapping strategy to perform current-clamp experiments identifying SNr neurons as DMS- or DLS-targeted based on the presence of an IPSP (Fig. 1D). We found that the spontaneous firing rate of DLS-targeted neurons (24.25 ± 2.5 Hz; 5/26 cells not spontaneously active) was significantly higher compared to DMS-targeted SNr neurons (17.66 ± 1. 35 Hz; 3/24 cells not spontaneously active) (Fig. 1E). The R_in_ of DMS-targeted SNr neurons (469.9 ± 40.9 MΩ) was significantly higher than DLS-targeted neurons (278.6 ± 25.4 MΩ) (Fig. 1F), which might indicate higher excitability of DMS-targeted neurons. We found fast rebound firing responses to hyperpolarization in most DMS-targeted and DLS-targeted neurons (Fig. 1G; fig. supp. F). However, a significantly higher number of DMS-targeted neurons displayed prolonged rebound firing responses when compared to DLS-targeted neurons (Fig. 1G; fig. supp. 1F). These results indicate that DMS-targeted and DLS-targeted SNr neurons differ in their intrinsic properties and post-inhibitory rebound responses.

We next studied the firing responses of SNr neurons during DMS or DLS terminal stimulation using trains of stimuli delivered at 10 and 20 Hz. We found that 10 Hz stimulation induced partial inhibition of firing at DMS-SNr DMS-^M^SNr (Light off: 11.2 ± 1.8 Hz; Light on 6.7 ± 1.5 Hz) and DLS-^L^SNr inputs (Light off: 19.8 ± 2.7 Hz; Light on 10 ± 2.5Hz), whereas a 20 Hz stimulation strongly inhibited firing in both the DMS-^M^SNr input (Light off: 10.6 ± 1.1 Hz; Light on: 0.9 ± 0.6 Hz) and the DLS-^L^SNr input (Light off: 15.2 ± 1.8 Hz; Light on 0.75 ± 0.72 Hz) (Fig. 1H). Post-inhibitory prolonged rebound responses emerged in most DMS-targeted and a subset of DLS-targeted SNr neurons in response to 10 or 20 Hz stimulation (Fig. 1I; Fig. supp. 1G-1H). These findings demonstrate that striatal inputs can induce post-inhibitory rebound firing in SNr neurons in a subpopulation-dependent manner.

### 2. Differential organization of STN inputs to the medial and lateral SNr

We next sought to determine the functional organization of STN inputs to the medial and lateral SNr. We virally expressed a Cre-dependent version of Chronos (AAV5-hSyn-FLEX-Chronos-GFP) in the STN of VGlut2-Cre mice (Fig. 2A). Here, we could not define ^M^SNr and ^L^SNr neurons based on their striatal input, but we based our classification on their spatial location. To ensure that we were recording from distinct SNr populations, we first confirmed that R_in_ was significantly higher in ^M^SNr than ^L^SNr neurons (Fig. supp. 2A), which matched previous data based on striatal input specificity (Fig. 1A). The amplitude of oEPSCs was significantly higher at STN-^L^SNr (0.52 ± 0.1 nA) over STN-^M^SNr (0.23 ± 0.05 nA) synapses (Fig. 2B), indicating that STN inputs are biased to the lateral SNr. We then reasoned that STN inputs might be scaled to the different intrinsic properties of SNr subpopulations. We found a significant correlation (r^2^ = 0.5338) between the amplitude of oEPSCs at STN-SNr synapses and the R_in_ of recorded neurons (Fig. 2C). Using paired stimuli delivered with a 50 ms interpulse interval, we found that the PPR at STN-^M^SNr synapses was facilitating (PPR: 1.3 ± 0.11) whereas no net facilitation or depression emerged at STN-^L^SNr (PPR: 1.04 ± 0.04) inputs, and that PPRs at STN-^M^SNr and STN-^L^SNr synapses were significantly different (Fig. 2D). Hence, subthalamic inputs are biased to the ^L^SNr, scaled to the R_in_ of their postsynaptic targets and higher release probability at STN-^L^SNr contributes to the target-specific organization of STN inputs.

**Figure 2:**
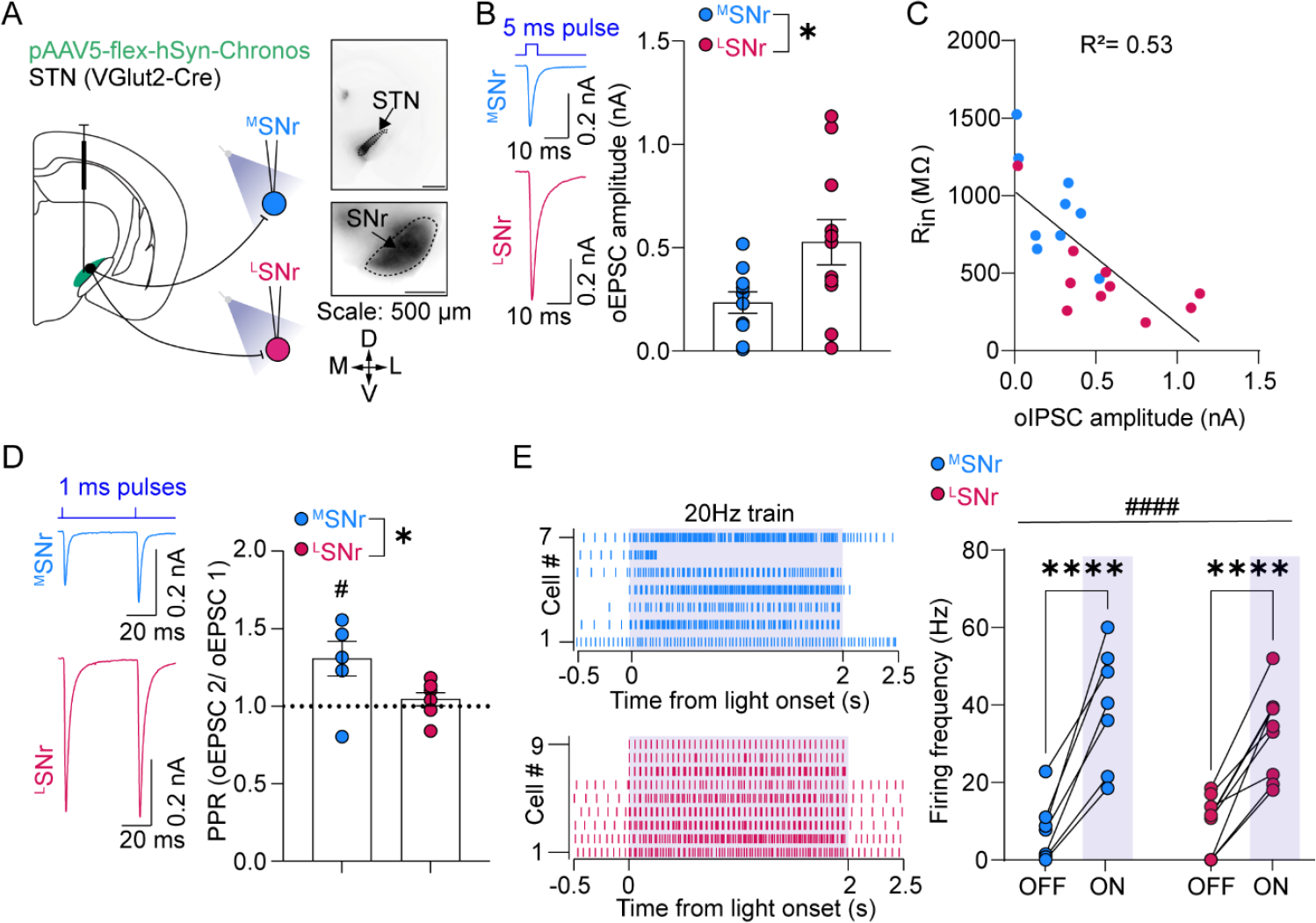
Differential organization of STN inputs to the medial and lateral SNr. **A**: Left: Viral injection schematic; Right: validation of Chronos-GFP expression in STN (injection site, top) and SNr (terminals, bottom). **B**: Amplitude of oEPSCs at STN-^M^SNr (N = 10 cells, 10 slices, 5 mice) and STN-^L^SNr (N = 11 cells, 10 slices, 6 mice) synapses (unpaired t-test, two-tailed, p = 0.03). **C:** Correlation of oEPSC amplitude and R_in_ at STN-SNr synapses (N = 19 cells, 14 slices, 6 mice) (R^2^ = 0.53; p = 0.0004). **D**: Paired pulse ratio at STN-^M^SNr (N = 6 cells, 6 slices, 5 mice) and STN-^L^SNr synapses (N = 7 cells, 6 slices, 4 mice) (one sample t-test hypothetical value 1, STN-^M^SNr p = 0.04, STN-^L^SNr p = 0.3; unpaired t-test, two-tailed, p = 0.04). **E**: Effects of STN inputs on ^M^SNr neuron firing (N = 7 cells, 7 slices, 4 mice) and ^L^SNr neuron firing (N = 9 cells, 8 slices, 3 mice) (2-way RM ANOVA, main effect of light F_1,14_ = 109.4, p < 0.0001; main effect of location F_1,14_ = 109.4 = 0.3, p = 0.14; Šídák’s multiple comparisons test ^M^SNr p < 0.0001; ^L^SNr P < 0.0001).

We next studied SNr firing responses while stimulating STN inputs using 20 Hz stimulus trains. The increase in firing relative to baseline produced by STN terminal stimulation was similar in ^M^SNr (Light off: 7.5 ± 3 Hz; Light on: 39.5 ± 5 Hz) neurons compared to ^L^SNr (Light off: 9.2 ± 2.5 Hz; Light on: 33 ± 3.7 Hz) neurons (Fig. 3H). Hence, STN inputs have different synaptic strengths in the different SNr subregions but are sufficient to induce firing in ^M^SNr and ^L^SNr neurons.

**Figure 3:**
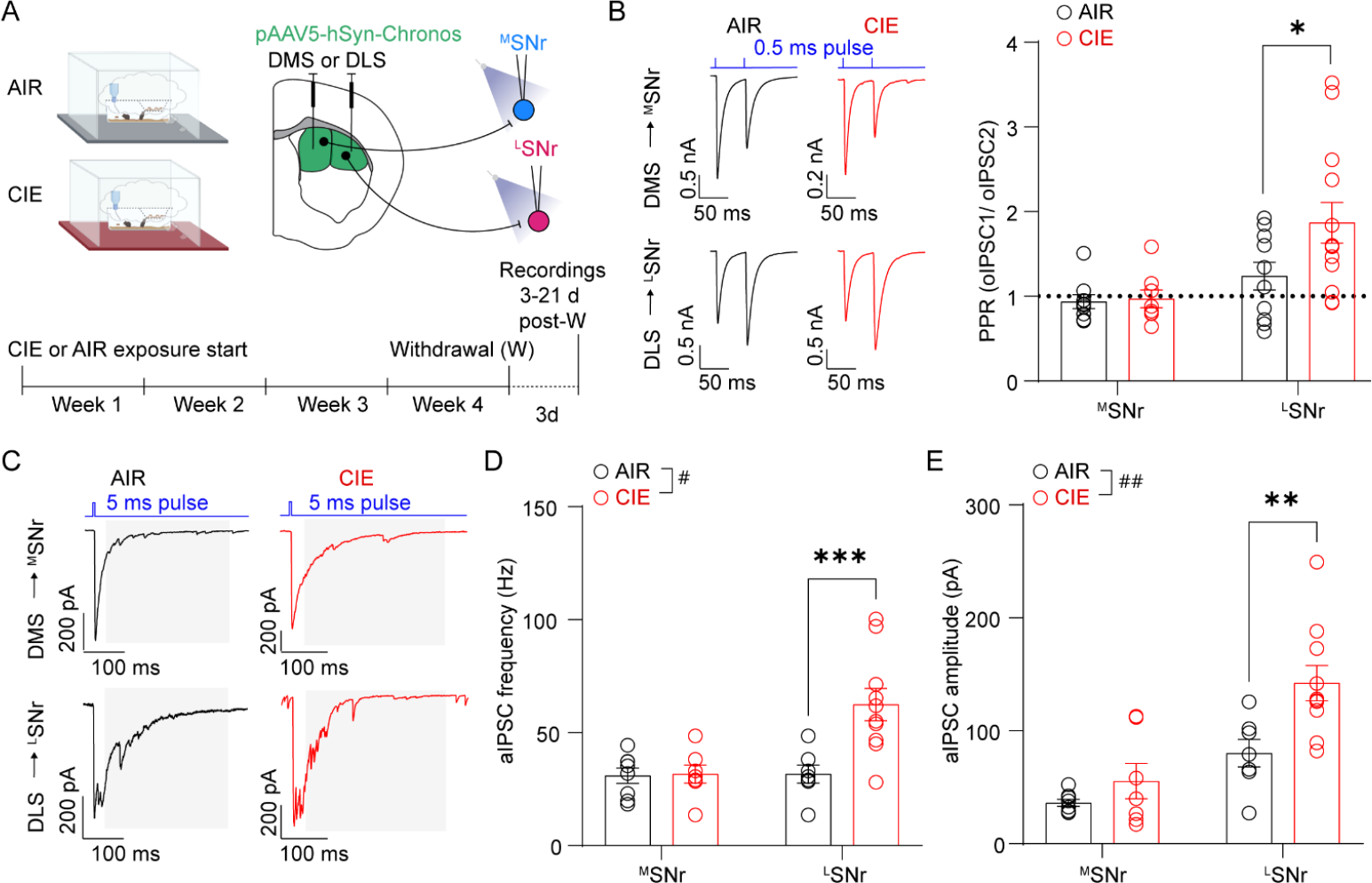
CIE potentiates DLS-SNr projections. **A**: Experimental schematic for viral injections and timeline of AIR or CIE exposure and electrophysiological recordings. **B**: CIE effects on paired pulse ratios at DMS-^M^SNr synapses (AIR: N = 9 cells, 7 slices, 4 mice; CIE: N = 7 cells, 7 slices, 5 mice) and DLS-^L^SNr synapses (AIR: N = 11 cells, 7 slices, 4 mice; CIE: N = 13 cells, 9 slices, 5 mice) (2-way RM ANOVA, main effect of group F_1, 36_ = 3.567, p = 0.067; main effect of synapse, F_1, 36_ = 8.758, p = 0.0054; Šídák’s multiple comparisons test ^M^SNr AIR vs ^M^SNr CIE p = 0.9783; ^L^SNr AIR vs ^L^SNr CIE p = 0.0178). **C**: Example traces of asynchronous IPSCs recorded at DMS-^M^SNr synapses and DLS-^L^SNr synapses in AIR and CIE groups. **D**: CIE effects on aIPSC frequency at DMS-^M^SNr (AIR: N = 8 cells, 5 slices, 3 mice; CIE: N = 7 cells, 4 slices, 3 mice) synapses and DLS-^L^SNr synapses (AIR: N = 7 cells, 4 slices, 3 mice; CIE: N = 10 cells, 9 slices, 5 mice) (2-way RM ANOVA, main effect of group, F_1,28_ = 8.314, p = 0.0075; main effect of synapse, F_1,28_ = 8.314, p = 0.0075; interaction group x synapse, F_1,28_ = 7.5, p = 0.0104; Šídák’s multiple comparisons test ^M^SNr AIR vs ^M^SNr CIE p = 0.99, ^L^SNr AIR vs ^L^SNr CIE p = 0.0007). **E**: CIE effects on aIPSC amplitude (AIR: N = 8 cells, 5 slices, 3 mice; CIE: N = 7 cells, 4 slices, 3 mice) at DMS-^M^SNr synapses and DLS-^L^SNr (AIR: N = 7 cells, 4 slices, 3 mice; CIE: N = 10 cells, 9 slices, 5 mice) synapses (2-way RM ANOVA, main effect of group, F_1,28_ = 9.254, p = 0.0051; main effect of synapse, F_1,28_ = 23.97, p < 0.0001; Šídák’s multiple comparisons test ^M^SNr AIR vs ^M^SNr CIE p = 0.55; ^L^SNr AIR vs ^L^SNr CIE p = 0.0044).

### 3. Altered DLS-SNr inputs but not DMS-SNr inputs following chronic alcohol exposure

Considering the previous findings describing alcohol effects on excitatory and inhibitory inputs to the DMS and DLS, we next asked whether chronic alcohol exposure changes DMS and DLS inputs to the SNr. We virally expressed Chronos in the DMS or DLS and subjected 3– 4-month-old C57BL mice to chronic intermittent alcohol vapor exposure (CIE) and a control group to air exposure (AIR) (Fig. 3A). We performed electrophysiological experiments during extended withdrawal (3-21 days post-withdrawal), a period associated with altered action control and cortico-striatal synaptic dysfunctions^25,26^ following CIE. We first investigated the presynaptic properties of DMS-SNr and DLS-SNr synapses between AIR and CIE mice. We found that CIE did not induce a change in PPR at DMS-SNr synapses, whereas the PPR at DLS-SNr synapses was significantly more facilitating (Fig. 3B). To investigate whether altered PPRs at DLS-SNr synapses were associated with changes in synaptic strength, we recorded asynchronous IPSCs (aIPSCs) (Fig. 3C). We found that CIE did not alter the frequency and amplitude of aIPSCs at DMS-SNr synapses (Fig. 3D, E). Conversely, we found that the amplitude and frequency of aIPSCs significantly increased at DLS-SNr synapses (Fig. 3D, E). Altogether, our results indicate that CIE alters presynaptic properties and increases synaptic strength of DLS-SNr synapses, but it does not affect the presynaptic properties and synaptic strength of DMS-SNr synapses.

### 4. STN inputs to the SNr are unaltered following chronic alcohol exposure

STN inputs to the SNr, particularly those to the lateral SNr, play a crucial role in action suppression^34^. Therefore, we investigated whether plasticity at STN-SNr inputs following chronic alcohol exposure might counteract or exacerbate imbalanced DLS-SNr synaptic inputs. VgluT2-Cre mice were injected with a Cre-dependent version of Chronos in the STN and allocated to AIR or CIE groups. Patch-clamp recordings were performed during prolonged withdrawal (3-21 days post-withdrawal) (Fig. 4A). We found that CIE did not affect the oEPSC amplitude at STN-^M^SNr and STN-^L^SNr synapses (Fig. 4B). Similarly, we found that CIE did not affect the PPR at STN-^M^SNr and STN-^L^SNr synapses (Fig. 4C). These results support the absence of CIE-induced changes in synaptic strength and presynaptic properties of STN inputs to the ^M^SNr and ^L^SNr.

**Figure 4:**
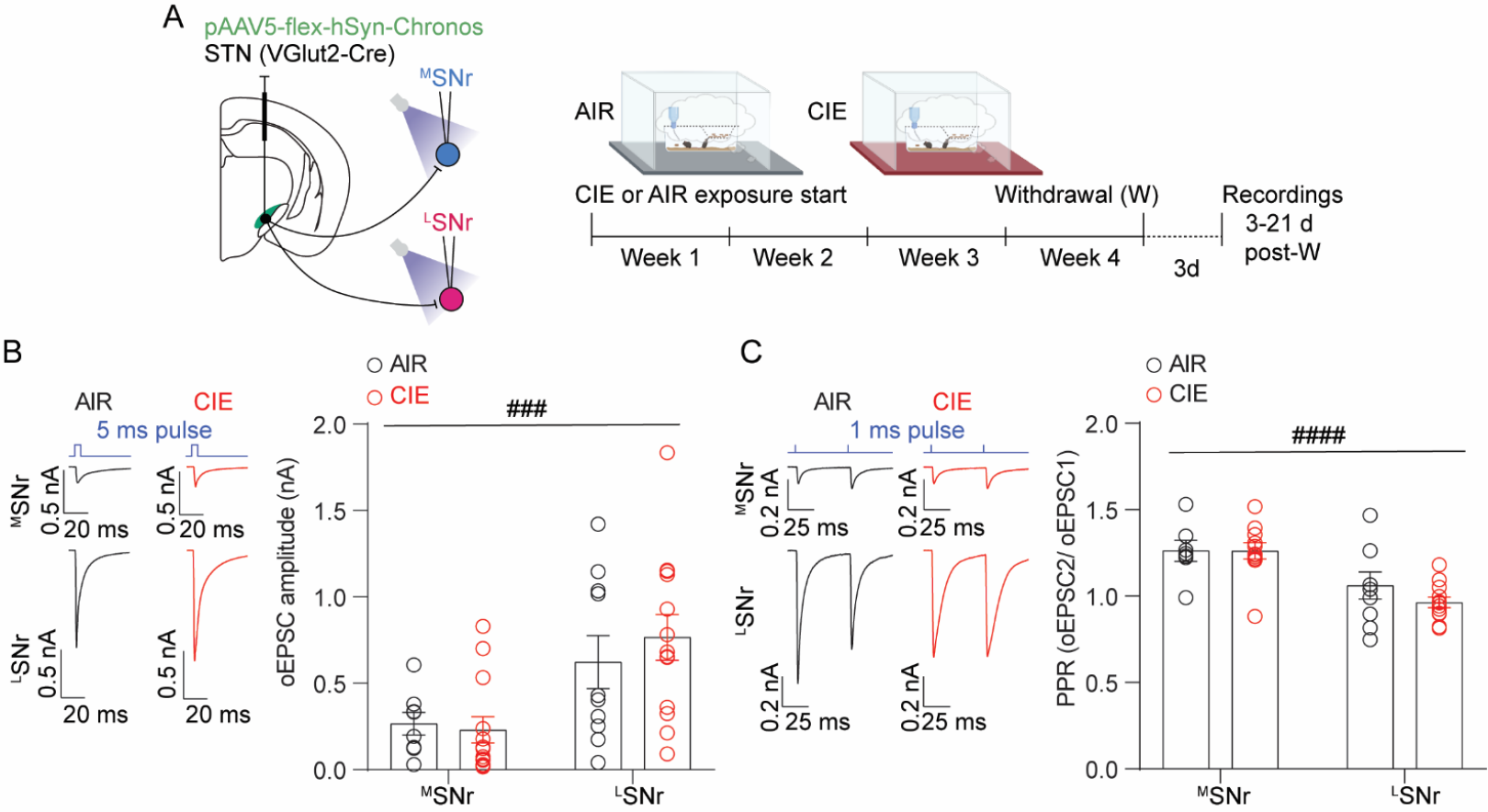
CIE does not affect STN-SNr inputs. **A**: Experimental schematic for viral injections and timeline of AIR or CIE exposure and electrophysiological recordings. **B**: CIE effects on the amplitude of oEPSCs at STN-^M^SNr (AIR: N = 8 cells, 8 slices, 3 mice; CIE: N = 13 cells, 13 slices, 4 mice) and STN-^L^SNr (AIR: N = 10 cells, 10 slices, 4 mice; CIE: N = 13 cells, 13 slices, 5 mice) synapses (2-way ANOVA, main effect of location, F_1, 40_ = 14.01, p = 0.0006). **C**: CIE effects on paired pulse ratios at STN-^M^SNr (AIR: N = 8 cells, 8 slices, 4 mice; CIE: N = 11 cells, 11 slices, 4 mice) and STN-^L^SNr (AIR: N = 9 cells, 9 slices, 4 mice; CIE: N = 13 cells, 13 slices, 4 mice) synapses (2-way ANOVA, main effect of location, F_1, 35_ = 21.20, p < 0.0001).

**Figure 5:**
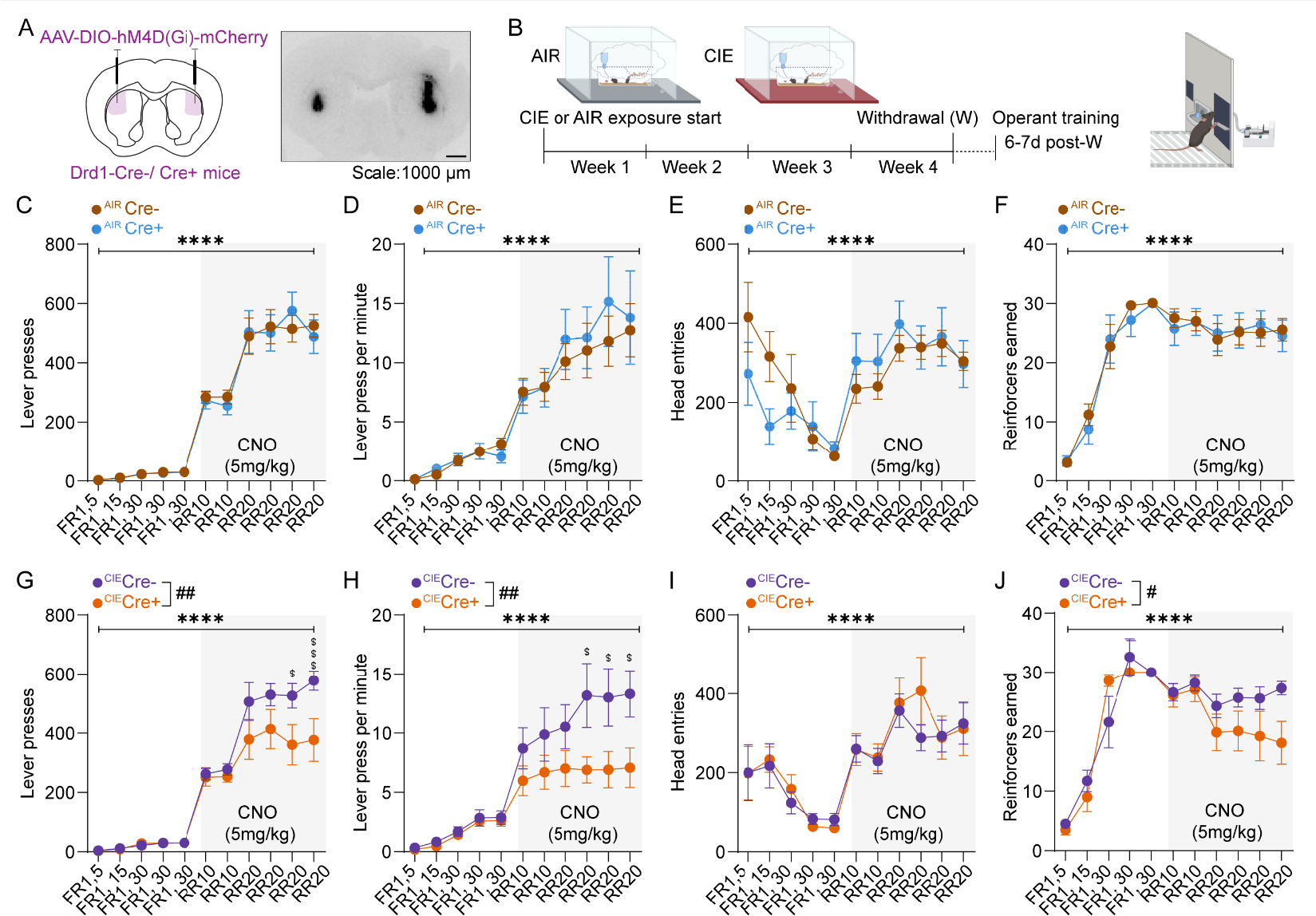
CIE induces a gain of function of DLS dSPNs for action control. **A**: Validation of viral expression of inhibitory DREADDs in the DLS. **B**: Timeline of CIE exposure and behavioral experiments. **C**: Lever presses in ^AIR^Cre-(N = 12) and ^AIR^Cre+ (N = 9) mice (2-way RM ANOVA, main effect of session, F_10, 190_ = 120.4, p < 0.0001). **D**: Lever presses per minute in ^AIR^Cre-(N = 12) and ^AIR^Cre+ (N = 9) mice (2-way RM ANOVA, main effect of session, F_10, 190_ = 28.43, p < 0.0001). **E**: Head entries in ^AIR^Cre-(N = 12) and ^AIR^Cre+ (N = 9) mice (Mixed-effect analysis, main effect of session, F_10, 189_ = 8.602, p < 0.0001). **F**: Reinforcers earned in ^AIR^Cre-(N = 12) and ^AIR^Cre+ (N = 9) mice (2-way RM ANOVA, main effect of session, F_10, 190_ = 42.76, p < 0.0001). **G**: Lever presses in ^CIE^Cre-(N = 10) and ^CIE^Cre+ (N = 8) mice (2-way RM ANOVA, main effect of session, F_10, 160_ = 87.82, p < 0.0001; main effect of genotype, F_10, 16_ = 4.88, p = 0.0421; interaction session X genotype, F_10, 160_ = 3.08, p = 0.0013; Šídák’s multiple comparisons test RR20-3 p = 0.0122; RR20-4 p = 0.0009). **H**: Lever presses per minute in ^CIE^Cre-(N = 10) and ^CIE^Cre+ (N = 8) mice (2-way RM ANOVA, main effect of session, F_10, 160_ = 24.8, p < 0.0001; main effect of genotype, F_10, 16_ = 3.672, p = 0.0734; interaction session X genotype, F_10, 160_ = 2.518, p = 0.0077; Šídák’s multiple comparisons test RR20-2 p = 0.0333; RR20-3 p = 0.0433; RR20-4 p = 0.0359). **I**: Head entries in ^CIE^Cre-(N = 10) and ^CIE^Cre+ (N = 8) mice (2-way RM ANOVA, main effect of session, F _10,160_ = 11.54, p < 0.0001). **J**: Reinforcers earned in ^CIE^Cre-(N = 10) and ^CIE^Cre+ (N = 8) mice (2-way RM ANOVA, main effect of session, F _10, 160_ = 31.54, p < 0.0001; interaction session x genotype F_1, 16_ = 2.087, p = 0.0283).

### 5. Chronic alcohol exposure shifts action control to direct pathway SPNs in the DLS

We used an operant conditioning task to determine how potentiated DLS-SNr inputs following CIE might impact action control. Drd1-Cre+ mice or Cre-littermates were injected with AAV-DIO-hM4D(Gi)-mCherry in the DLS and randomly allocated to AIR or CIE groups (Fig. 4A). 6-7 days after withdrawal, AIR (Cre-, Cre+) and CIE (Cre-, Cre+) mice underwent operant conditioning for 20% sucrose reward that began with 5 days of fixed ratio 1 training and then progressed to 2 days of random ratio 10 and 4 days of random ratio 20 training (Fig. 4B). Mice received CNO (5 mg/kg) injections during random ratio (RR) training to inhibit dSPNs in the DLS. In AIR mice, chemogenetic inhibition of DLS-dSPNs did not affect the behavioral performance measured as number of lever presses (Fig. 4C), frequency of lever presses (Fig. 4D), head entries (Fig. 4E) and reinforcers earned (Fig. 4F). These results indicate that the function of DLS dSPNs was not necessary to sustain action execution in AIR mice.

In CIE mice, chemogenetic inhibition of DLS-dSPNs significantly reduced the total number of lever presses (Fig. 4G), the lever pressing frequency (Fig. 4H) and the number of reinforcers earned (Fig. 4J), whereas the number of head entries remained unchanged (Fig. 4I). This analysis indicated that hM4D(Gi)-expressing ^CIE^Cre+ mice showed fewer and slower lever presses and fewer rewards earned during CNO treatment than ^CIE^Cre-mice. As opposed to control mice, these results indicate that the function of DLS dSPNs becomes necessary for action execution in CIE mice. Our results reveal a gain of function in dSPNs in the DLS for action control following chronic alcohol exposure.

## Discussion

Here we describe a distinctive functional organization of striatal and subthalamic synaptic inputs to the medial and lateral substantia nigra pars reticulata and investigate their vulnerability to the effects of chronic alcohol exposure. We find that inputs from the dorsolateral striatum are potentiated following chronic inhalational alcohol exposure, likely contributing to altered basal ganglia-mediated action control following this exposure.

### On the synaptic organization of the SNr

Our findings indicate that the synaptic properties of striatal and subthalamic synaptic inputs to the SNr are specialized depending on the targeted neuronal subpopulation. The subpopulations of the SNr have been previously subdivided based on their source of striatal input, output connectivity, and molecular identity. We propose that specialized synaptic input properties contribute to fine-tuning signal processing in parallel SNr subcircuits. We find that DLS projection neurons primarily synapse onto parvalbumin-expressing (PV^+^) SNr neurons, whereas the DMS neurons primarily innervate non-parvalbumin-expressing (PV^-^) SNr neurons. Our results agree with an enrichment of PV^+^ neurons in the ventrolateral SNr^24,35^. Previous studies indicate that the second largest subpopulation of SNr neurons consists of GAD2^+^/VGAT^+^/PV^-^ neurons^13^ or GAD2^+^ neurons^35^, and future studies will address whether DMS-targeted SNr neurons fall within this category. PV^+^-expressing and PV^-^ -non-expressing subpopulations of SNr neurons have been previously suggested to control distinct aspects of motor performance^23^ and sleep states^35^. Our work directly links molecularly identified SNr neurons to their striatal inputs, supporting a model whereby the DMS and DLS support distinct functional outputs through molecularly distinct subpopulations of SNr neurons.

How are striatal inputs onto subpopulations of SNr neurons organized? We reveal that DMS-SNr and DLS-SNr inputs differ in their presynaptic properties. Previous reports using electrical or optogenetic stimulation found that short-term facilitation is characteristic of striato-nigral synapses^36,37^. Here, we observe short-term facilitation at DLS-SNr synapses. Conversely, we find short-term depression at DMS-SNr synapses. These distinct synaptic properties may reflect distinct functional demands of postsynaptic neurons. Here we show that DMS-targeted neurons exhibit higher R_in_ than DLS-targeted SNr neurons. We find higher baseline firing rates in DLS-targeted over DMS-targeted SNr neurons. Our results support previous findings indicating a medio-lateral gradient of intrinsic properties within the SNr, whereby medial neurons have higher R_in_ and lower firing rates^13^. Similarly, our results agree with higher R_in_ and lower baseline firing rate in PV^-^ over PV^+^ SNr neurons^24^. Synaptic facilitation may be required at DLS-SNr synapses to sustain postsynaptic responses in SNr neurons. In contrast, synaptic depression at DMS-SNr synapses may contribute to faster responses in the highly excitable DMS-targeted neurons. The differential presynaptic properties of DMS and DLS synapses could also contribute to tuning specific firing responses in SNr neurons with distinct postsynaptic targets^13,21^. In particular, subpopulations of SNr neurons have been indicated to differ in their rebound firing capability, the lowest being dorsal-raphe projecting neurons and the highest medulla-projecting neurons^13^. Using negative current injections, we find that both DMS-targeted and DLS-targeted SNr neurons exhibit fast rebound responses, and that a significantly higher number of DMS-targeted neurons display prolonged rebound responses. We demonstrate that striatal inputs are capable of eliciting rebound responses in SNr neurons. At DMS-SNr inputs, most targeted neurons exhibit prolonged rebound, whereas at DLS-SNr inputs we find < 50% of neurons displaying prolonged rebound. Our results support a circuit logic whereby DMS and DLS synaptic inputs are differentially organized to allow specific firing responses in subpopulations of SNr neurons. These findings may generalize to other striatonigral projections, including the one from the ventrolateral striatum (VLS)^12,16^.

Our results also indicate a subregion-specific organization of indirect pathway STN inputs. We find that subthalamic synaptic inputs have higher strength in the lateral compared to medial SNr. We also find that a higher release probability characterizes STN inputs to the lateral SNr than STN inputs to the medial SNr. Furthermore, we find that STN inputs scale with the R_in_ of SNr neurons, thus being capable of eliciting firing to a similar extent in both medial and lateral SNr neurons. The STN inputs to the medial and lateral SNr may arise from topographically parallel projections and spatio-molecularly distinct subpopulations^38^. Collectively, our findings support a model in which synaptic properties and intrinsic SNr neuron properties interplay to allow for signals to be processed according to the different demands imposed by the distinct target regions and functional outputs of medial and lateral subpopulations of SNr neurons.

We did not directly investigate the mechanisms driving the functional differentiation of direct and indirect pathway synaptic inputs to distinct SNr subpopulations. We propose that input cell type differences across subcircuits and differences in presynaptic neuromodulation influence the distinctive synaptic organization of striato-SNr and subthalamic-SNr subcircuits. The subpopulation-specific synaptic organization we revealed might generalize to other synaptic inputs to the SNr, including the pallidonigral projection. We speculate that a consequence of this differential organization might be a differential vulnerability of synaptic inputs to the SNr in pathological conditions affecting basal ganglia functioning, including Alcohol Use Disorder and Parkinson’s Disease^26,39,40^.

### Selective vulnerability of DLS-SNr inputs underlies altered action control following chronic alcohol exposure

Chronic alcohol exposure induces widespread adaptations in the dorsal striatum, but the direction of these changes is specific to different DS subregions and cell types, thereby affecting distinct behavioral outputs^30–33,42,43^. We found that chronic alcohol exposure potentiates DLS-SNr synapses, leaving DMS-SNr synapses unchanged. This form of synaptic potentiation is accompanied by a gain of function of DLS dSPNs for action control following chronic alcohol exposure.

Previous studies demonstrated that chronic alcohol exposure engages several circuit mechanisms to disinhibit DLS output. These include decreased GABAergic synaptic transmission onto SPNs in DLS in a mouse model of binge drinking^43^. Similarly, chronic alcohol drinking reduces GABAergic synaptic inputs, increases glutamatergic inputs, and increases the excitability of SPNs in the putamen of macaque monkeys^44,45^. We previously reported that chronic intermittent ethanol vapor exposure promotes dendritic arborization of DLS SPNs ^46^. Chronic alcohol exposure disinhibits cortical inputs to the DLS through reduced endocannabinoid-dependent plasticity^46^. Potentiated anterior insular cortex inputs to the DLS maintain alcohol binge drinking in male mice^42^, due to impaired µ-opioid dependent long-term depression^47^. These studies collectively revealed a disinhibition of DLS neurons following acute and chronic alcohol exposure. However, the studies did not discriminate between effects on direct and indirect pathway striatal output. Our study indicates a selective vulnerability of dSPN-SNr DLS inputs to chronic alcohol exposure. We found that the PPR of IPSC at DLS-SNr inputs was significantly more facilitating following chronic alcohol exposure, indicating decreased presynaptic probability of GABA release following chronic alcohol exposure. Therefore, the increased frequency and amplitude of asynchronous IPSCs might emerge due to a postsynaptic mechanism. Future studies will investigate whether this results from an increased number of GABAergic synapses, an increased number of GABA receptors, or changes in channel conductance due to modifications in their subunit composition (^3,4^).

A prominent circuit model posits that weakening the associative OFC-DMS^32,33^ pathway mediates alcohol-induced loss of goal-directed behavior. In this regard, the absence of alcohol-induced maladaptations at downstream DMS-medial SNr synapses indicates that alcohol-induced disinhibition of DMS dSPNs^30,31,49,50^ might primarily affect striatal integration of cortical and thalamic inputs. On the contrary, our behavioral and electrophysiology data support an alcohol-induced sensorimotor over-engagement (DLS-lateral SNr) during action execution, corroborating previous findings on heightened DLS neuron function during visual discrimination^46^. Finally, considering their role in canceling actions^34^ and the lack of alcohol-induced changes at STN-SNr inputs, our data further support a model where alcohol exposure confers to DLS-SNr an advantage over STN-SNr input in determining lateral SNr output and action execution.

### Materials and methods

#### Ethical approval

Experiments were conducted in accordance with the National Institutes of Health’s Guidelines for Animal Care and Use, and all experimental procedures were approved by the National Institute on Alcohol Abuse and Alcoholism Animal Care and Use Committee (protocol # LIN-DL-1). Mice were housed in groups of 2-4 in the NIAAA vivarium on a 12-hour light cycle with ad libitum access to food and water.

#### Animals

VGluT2-IRES-Cre mice (B6J.129S6(FVB)-Slc17a6tm2(cre)Lowl/MwarJ, Cat. No. JAX 028863), PV-IRES-Cre mice (B6.129P2-Pvalbtm1(cre)Arbr/J, Cat. No. JAX 017320), Td-Tomato mice (B6-Cg-Gt(ROSA)26Sortm14(CAGtdTomato)Hze/, Cat. No. 007914) and C57BL-6J mice were purchased from the Jackson laboratory (JAX; Bar Harbor, ME, USA). Heterozygous Drd1-Cre (STOCK Tg(Drd1-cre)FK150Gsat/Mmucd) mice were purchased from Gensat and crossed with C57BL-6J mice. VGluT2-IRES Cre and PV-IRES-Cre homozygous and heterozygous (het) male or female mice were crossed with C57BL-6J. Homozygous PV-IRES-Cre mice were crossed with homozygous Td-Tomato mice to generate PVCre-TdTom mice. Experiments were conducted on male and female mice aged >3 months. All mice involved in the CIE experiments initiated the CIE exposure between 3 and 4 months of age. Hence, mice were tested for behavior or slice electrophysiology between 4 and 5 months of age. Genotyping was performed by polymerase chain reaction (PCR) on genomic DNA from ear biopsies.

#### Surgical procedures

Deep anesthesia was performed in an induction chamber using 5% Isoflurane. Animals were quickly transferred to a stereotaxic frame that delivered isoflurane at 1-3% through the mouthpiece for the whole duration of the surgery. A Hamilton syringe pre-loaded with the viral solution was inserted in the brain parenchyma to target the DLS (AP: +0.9; ML: ±2.4; DV: -3.2), DMS (AP: +0.9; ML: ±1.4; DV: -3.2) or STN (AP: -1.9; ML: ±1.7; DV: - 4.5). 200 nL of pAAV5-Syn-Chronos-GFP (Addgene, plasmid #59170, 5×10^12^ vg/mL) were injected in the DMS or DLS (injection rate 25 nl/ min) of C57BL-6J mice. 300 nL of AAV5-hSyn-FLEX-Chronos-GFP (UNC viral core, 7×10^12^ vg/mL) were injected in the STN (injection rate 50 nL/ min) of VGlut2-IRES-Cre mice. 300 nL of AAV2-hSyn-DIO-HM4D(Gi)-mCherry (Addgene, plasmid #50475, 7×10^12^ vg/mL) were injected in the DLS of Drd1-Cre+ mice and their Cre-littermates. Following the injection of the viral solution, the syringe was kept in place for 5 minutes and then retracted. The wounded skin was closed using skin glue (Vetbond, 3M). Post-operative care included the administration of ketoprofen (5mg/ kg, s.c.) following surgeries and two days of recovery, where half of the cage was placed on a heated pad.

#### Brain slice preparation for patch-clamp experiments and validation of viral expression

Mice were deeply anesthetized with isoflurane and decapitated. The brain was collected and transferred to a slicing chamber filled with ice-cold sucrose-based cutting solution containing the following (in mM): 194 sucrose, 30 NaCl, 4.5 KCl, 26 NaHCO_3_, 1.2 NaH_2_PO_4_, 10 D-glucose, 1 MgCl_2_, saturated with 95% O_2_/5% CO_2_. Coronal sections (250 μm) containing the SNr, STN, or striatum were obtained using a Leica VT1200S Vibratome (Leica Microsystems, Buffalo Grove, IL) and transferred to an incubation chamber filled with artificial cerebrospinal fluid (aCSF) containing (in mM): 124 NaCl, 4.5 KCl, 26 NaHCO_3_, 1.2 NaH_2_PO_4_, 10 D-glucose, 1MgCl_2_, and 2 CaCl_2_, saturated with 95% O_2_/5% CO_2_). Slices were incubated for 45-60 minutes at 32°C and then maintained at room temperature. Slices containing the STN, SNr, or striatum were transferred to an aCSF-filled petri dish, and low-magnification epifluorescence images were acquired using a Zeiss Axiozoom microscope (Carl Zeiss, Oberkochen, Germany) via a Zeiss Axiocam MR monochrome CCD camera for validation of viral expression.

#### Whole-cell patch-clamp recordings and analysis

Hemislices were transferred to a recording chamber perfused with aCSF at 30–32 °C. Neurons were visualized with an upright microscope (Model BX51WI, Olympus, Waltham, MA) using a 10x air objective or 40x water objective (LUMPlanFL, 0.80 NA) and connected to a CCD camera (SciCam Pro, Scientifica) controlled via the Ocular Imaging acquisition software (Teledyne Photometrics, Tucson, Arizona). Images were obtained using the 10x objective to track the approximate location of the recorded neurons. Recordings from SNr neurons were obtained using micropipettes (2–4 MΩ) made from 1.5 mm OD/1.12 mm ID borosilicate glass with a filament (World Precision Instruments, Sarasota, FL) pulled with a P-97 puller (Sutter Instruments, Novato, CA). The intracellular solution for voltage clamp recordings of inhibitory post-synaptic currents (IPSCs) contained the following (in mM): 150 CsCl, 10 HEPES, 2.0 MgCl_2_, 0.3 Na-GTP, 3.0 Mg-ATP, 0.2 BAPTA-4Cs, 5 QX-314. The intracellular solution for voltage clamp recordings of excitatory post-synaptic currents (EPSCs) contained the following (in mM): 114 CsMeSO_3_, 5.0 NaCl, 1.0 TEA-Cl, 10 HEPES, 5.0 QX-314, 1.1 EGTA, 0.3 Na-GTP, 4.0 Mg-ATP. The intracellular solution for current clamp recordings contained the following (in mM): 140 mM K-Glu, 10mM HEPES, 0.1 mM CaCl_2_, 2 mM MgCl_2_, 1 mM EGTA, 2 mM ATP-Mg, 0.2 mM GTP-Na. Voltage clamp recordings were performed using Multiclamp 700B and a Digidata 1550B digitizer using a low-pass filter of 2 kHz and a sampling frequency of 10 kHz. Current clamp recordings were performed using Multiclamp 700B and a Digidata 1550B digitizer using a low-pass filter of 2-10 kHz and a sampling frequency of 10 kHz. Data were acquired and analyzed using pClamp 10.3 or pClamp 10.6 software (Molecular Devices, Sunnyvale, CA). To isolate IPSCs, the AMPA receptor antagonist DNQX (10 µM) and the NMDA receptor antagonist DL-AP5 (50 µM) were continuously bath applied. To isolate EPSCs, the GABAA antagonist Gabazine (10 µM) was continuously bath-applied. To record asynchronous IPSCs, a modified aCSF was used, whereby Ca^2+^ was replaced by Sr^2+^ 4mM.

SNr neurons were characterized in voltage clamp experiments (holding voltage: -50 mV) for their location, appearance, and firing frequency assessed in tight-seal (resistance >500 MΩ) cell-attached mode before the break-in, and absence of a hyperpolarization-activated current (Ih) measured using a -50 mV step in voltage-clamp mode after break-in^37^. In experiments conducted in PVCre-TdTom mice, the following procedure was followed: neurons were visualized using DIC and a tight-seal was established; next, yellow light (509-551 nm) was briefly delivered via field illumination through the microscope objective (generated via an X-Cite LED driver, Lumen dynamics, and filtered through a Cy3 filter) to label neurons as TdTom+ or TdTom-; following breakthrough, connectivity was assessed; the field of view of the slice was then changed to reduce sampling bias in the next recording attempt.

oIPSC and oEPSC amplitudes, latency to peak, PPRs, and input resistance in voltage clamp experiments (measured from the steady state of a -50 mV step) were analyzed using clampfit (pClamp suite). Input resistance in current clamp experiments (measured from the steady state of a - 50 pA, 500 ms current step) was analyzed using Easy Electrophysiology (Easy Electrophysiology Ltd, London, England). Asynchronous IPSC frequency and amplitude were analyzed using Easy Electrophysiology. A minimum of 6 trials were averaged to calculate oIPSC amplitudes, PPRs, aIPSC amplitude and frequency, and input effects on SNr neuron firing. For rebound firing analysis, a baseline was defined for each cell as the average firing frequency recorded 1 second before a -50 pA, 500 ms current step, or before optogenetic stimulation of striatal inputs. Rebound responses were then analyzed by normalizing to baseline the first instantaneous frequency (fast rebound) or the average firing frequency 400 ms (prolonged rebound) following negative current injections, or 1 second following optogenetic stimulation of striatal inputs. Neurons in which firing increased by more than 5% from to baseline were defined as “rebounders.”

#### Optogenetic stimulation protocols for patch-clamp recordings

The light was generated via an X-Cite LED driver (Lumen dynamics), delivered via field illumination through the microscope objective, and filtered through a single band filter to produce ∼470 nm illumination. To measure absolute amplitudes of oIPSCs and oEPSCs and evoked asynchronous IPSCs, blue light pulses of 5 ms and an optical power of ∼10 mW were used. To measure paired-pulse ratios, blue light pulses of 0.5-1 ms width with an inter-stimulus interval of 50 ms were used (the optical power was adjusted to the minimum required to produce a stable response, range 0.56 – 6 mW). For current-clamp experiments, 2-second-long trains of stimuli at 10 or 20Hz were delivered, using a pulse width of 1-5 ms and optical power of ∼5 mW. For all experiments, light pulses were delivered every 20-30 seconds.

#### Procedure for Chronic Intermittent ethanol exposure

Ethanol vapor exposure was conducted in Plexiglas chambers (Plas Labs Inc., Lansing, MI). A subset of chambers was used to expose mice to vaporized ethanol (CIE group), and a separate subset was used to expose mice to air (AIR group). The chambers were connected to a vaporizer. In the ethanol chamber, 95% ethanol was vaporized through air flow at a rate of 1.5-3 L/min. Vaporized ethanol was combined with another air stream to produce a total flow rate of ∼10 L/ min in each chamber. A similar rate of delivery was provided in the air chamber. The rate was adjusted throughout the cycle to produce vapor ethanol concentrations of 0.160-0.240 mg/L. Vapor ethanol concentrations were assessed using a breath analyzer. Mice were exposed to 4 cycles of CIE, each consisting of 4 consecutive days in which mice were exposed to 16 hours of ethanol vapor (CIE group) or air (AIR group) followed by 8 hours of withdrawal. In between cycles, mice underwent 72h of withdrawal. Mice were not given any loading dose of ethanol or pyrazole injections before the exposure. This procedure yielded average blood ethanol levels of 165.2 ± 10.6 mg/dl. Throughout the 4 weeks of CIE exposure, mice remained in the room containing the vapor chambers. This room had the same light cycle, temperature, and humidity settings as the colony room where mice were previously housed and where mice returned following CIE.

#### Procedure for blood ethanol concentration measurements

Blood was collected maximum twice from each mouse throughout the 4 weeks of CIE, with a typical interval of 2 weeks between collections. The collection was done through a caudal vein tail nick. Blood (<20 µL) was collected from the tail vein in heparinized hematocrit tubes (Fisherbrand, Cat. No. 22-362-566) and rapidly transferred for centrifugation at 10.000 rpm for 5-10 minutes. The serum was isolated, transferred to Eppendorf tubes, and diluted for further processing. Blood ethanol concentration was measured using a colorimetric assay (Pointe Scientific, MI, USA).

#### Operant conditioning

48 hours following the end of the 4^th^ CIE cycle, mice underwent food restriction to reach ∼90% of their body weight. Operant training started 6-7 days following the end of the 4^th^ CIE cycle. Mice received injections of saline or clozapine-N-oxide (CNO, 5mg/ kg) 1h before the start of each operant conditioning session. Mice were trained to lever press for 20% sucrose in operant boxes (MedAssociates). Training began with 1 day of magazine training in which no lever was presented, but sucrose was delivered at random intervals (average 60 seconds) in the magazine. All mice successfully learned to identify the magazine and consume the reward and were moved to the operant training, which started the following day. During initial training, mice were trained to lever press under a fixed ratio (FR) schedule (1 lever press = 1 reward) for 5 days, with a time criterion of 60 minutes, after which the session was interrupted (Day 2: 5 rewards; Day 3: 15 rewards; Day 4-6: 30 rewards). Mice were then moved to a random ratio (RR) schedule in which they received a reward after an average of 10 lever presses (RR10, Day 7-8) and then after an average of 20 lever presses (RR20, Day 9-12), for a maximum of 30 rewards over 60 minutes.

#### Immunohistochemistry and confocal imaging

Mice were anesthetized with pentobarbital (50 mg/ kg) before transcardial perfusion with 1x PBS, followed by 4% formaldehyde (FA) (4% paraformaldehyde in 1x PBS). Brains were kept in 4% FA overnight and then transferred to 1x PBS until slicing. Free-floating sections (50 µm) were cut using a Pelco easySlicer Vibratome (Ted Pella Inc., Redding, CA, USA). Slices were washed for 3 x 5 minutes in 1x PBS and then incubated in a blocking solution containing 5% normal goat serum in PBS-T (0.2% Triton X-100). Slices were then incubated over 2 nights in 1x PBS containing the primary antibody (Rabbit anti-RFP, Rockland, Cat. No 600-401-379). Slices were then washed 3 x 5 minutes in 1x PBS and incubated in 1x PBS containing the secondary antibodies for 2 hours (Alexa Fluor 594 goat anti-rabbit, Invitrogen A11012, 1:500). A final wash of 3 x 5 minutes was performed before mounting using DAPI Fluoromount-G® (Southern Biotech, Cat No. 0100-20). Low-magnification epifluorescence images were obtained through a Zeiss Axiozoom microscope (Carl Zeiss, Oberkochen, Germany) equipped with standard Cy3 filters via a Zeiss Axiocam MR monochrome CCD camera.

#### Viruses and Reagents

Viruses were aliquoted in 2.5-5 µL aliquots and stored at -80 until use. All drugs (DNQX, DL-AP5, Gabazine) were purchased from Tocris.

#### Statistical analysis

Statistical analysis was performed in GraphPad Prism (GraphPad Software, La Jolla, California), and data was pre-processed in Excel. The Shapiro-Wilk test was used to determine if data were normally distributed. A ROUT test (Q = 0.1%) was used to identify statistical outliers. Significance was set at p < 0.05 in all analyses and indicated in figures (^*,#,$^ for p < 0.05; ^**,##,$$^ for p < 0.01; ^***,###,$$$^ for p < 0.001; ^****,####,$$$$^ for p < 0.0001). Statistical significance is indicated in the figures and figure legends. All data are reported as mean + sem. Data were analyzed using two-way RM ANOVA or mixed-effects analysis with no correction. Post-hoc comparisons were analyzed using the Šidák post-hoc test where appropriate. All other paired comparisons, unpaired t-tests, or Mann-Whitney test were performed.

## Author contributions

Conceptualizations: G.S., S.B., D.M.L.; Methodology: G.S. and S.B.; Investigation: G.S., A.G. Supervision and funding acquisition: D.M.L.; Writing-original draft: G.S.; Writing-review and editing: S.B., D.M.L.

## Acknowledgments

The authors thank Dr. Pamela Ilse Alonso and Dr. Jeong Lee for assistance with CIE procedures and helpful discussions, Zev Jarret and Stephanie Ramos Maciel for support in surgical procedures and preparation of slices for validation of viral expression. The authors also thank the NIAAA animal care staff for supporting animal husbandry and veterinary care, and Dr. Guoxiang Luo for performing genotyping.

## Competing interests

The authors declare they have no competing interests.

## Funding

This work was supported by the National Institutes of Health, National Institute on Alcohol Abuse and Alcoholism, Division of Intramural Clinical and Biological Research (ZIAAA000416). SB was supported by the Center on Compulsive Behaviors at NIH.

---

**Figure supplementary 1:**
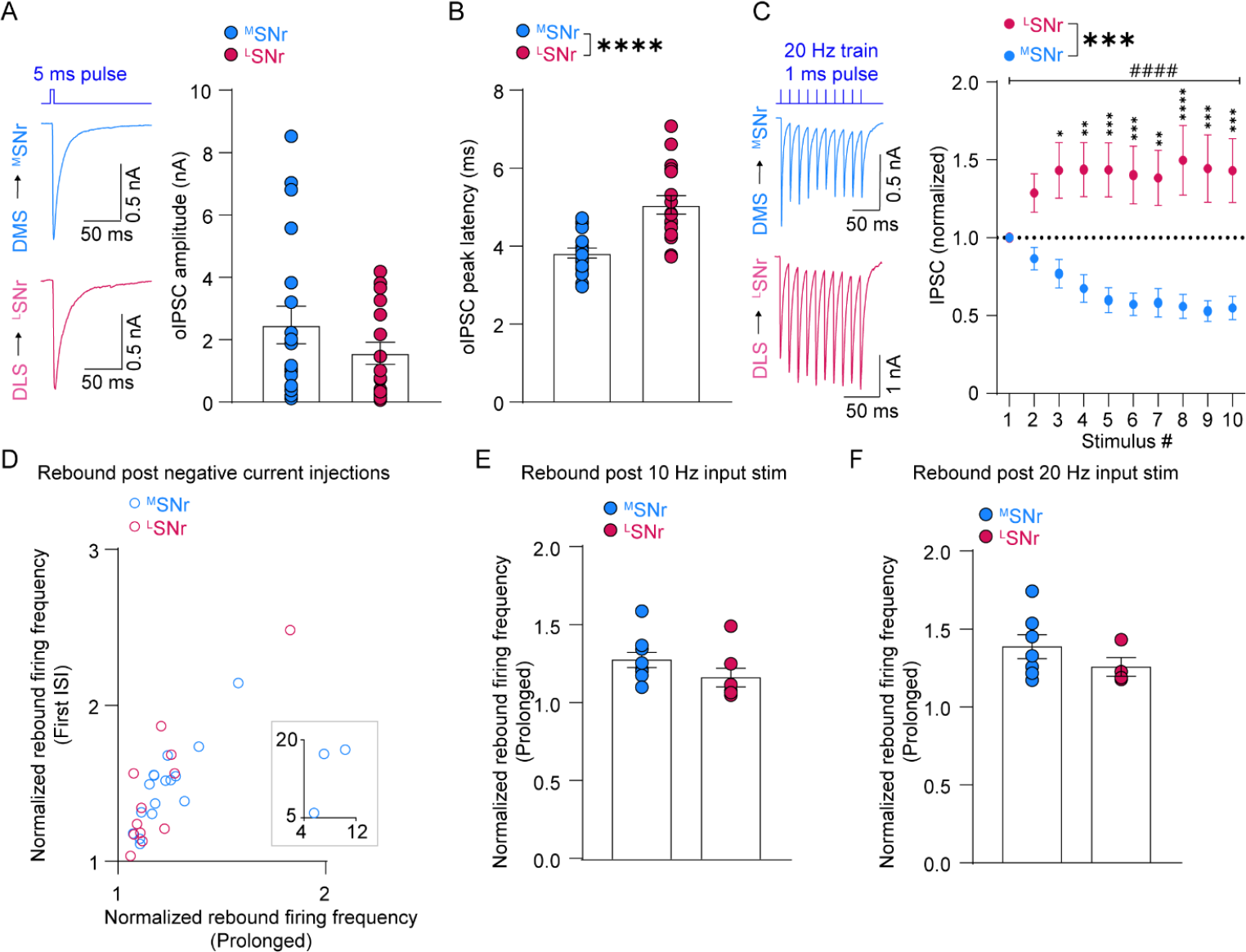
Synaptic characterization of DMS-SNr and DLS-SNr projections. **A**: Amplitude of oIPSCs at DMS-^M^SNr (N = 19 cells, 12 slices, 6 mice) and DLS-^L^SNr (N = 17 cells, 14 slices, 7 mice) synapses (Mann-Whitney test, two-tailed, p = 0.4). **B**: Latency to peak of oIPSCs at DMS-^M^SNr (N = 19 cells, 12 slices, 6 mice) and DLS-^L^SNr (N = 17 cells, 14 slices, 7 mice) synapses (unpaired t-test, two-tailed, p < 0.0001). **C**: oIPSCs (normalized to 1^st^) during a 20 Hz train of 10 pulses at DMS-^M^SNr (N = 10 cells, 7 slices, 5 mice) and DLS-^L^SNr synapses (N = 11 cells, 10 slices, 5 mice) (2-way RM ANOVA, main effect of synapse, F_1, 19_ = 14, p = 0.0014; interaction synapse x pulse, F_9, 171_ = 12.8, p < 0.0001; Šídák’s multiple comparisons tests). **D**: Amplitude (normalized to baseline firing rates) of fast (y axis) and prolonged (x axis) post-inhibitory rebound responses (fig. 1G) in ^M^SNr and ^L^SNr neurons; inset: high-rebounding ^M^SNr neurons. **E**: Amplitude of post-inhibitory rebound responses induced by stimulating DMS or DLS inputs using a 10 Hz train. **F**: Amplitude of post-inhibitory rebound responses induced by stimulating DMS or DLS inputs using a 20 Hz train.

**Figure supplementary 2:**
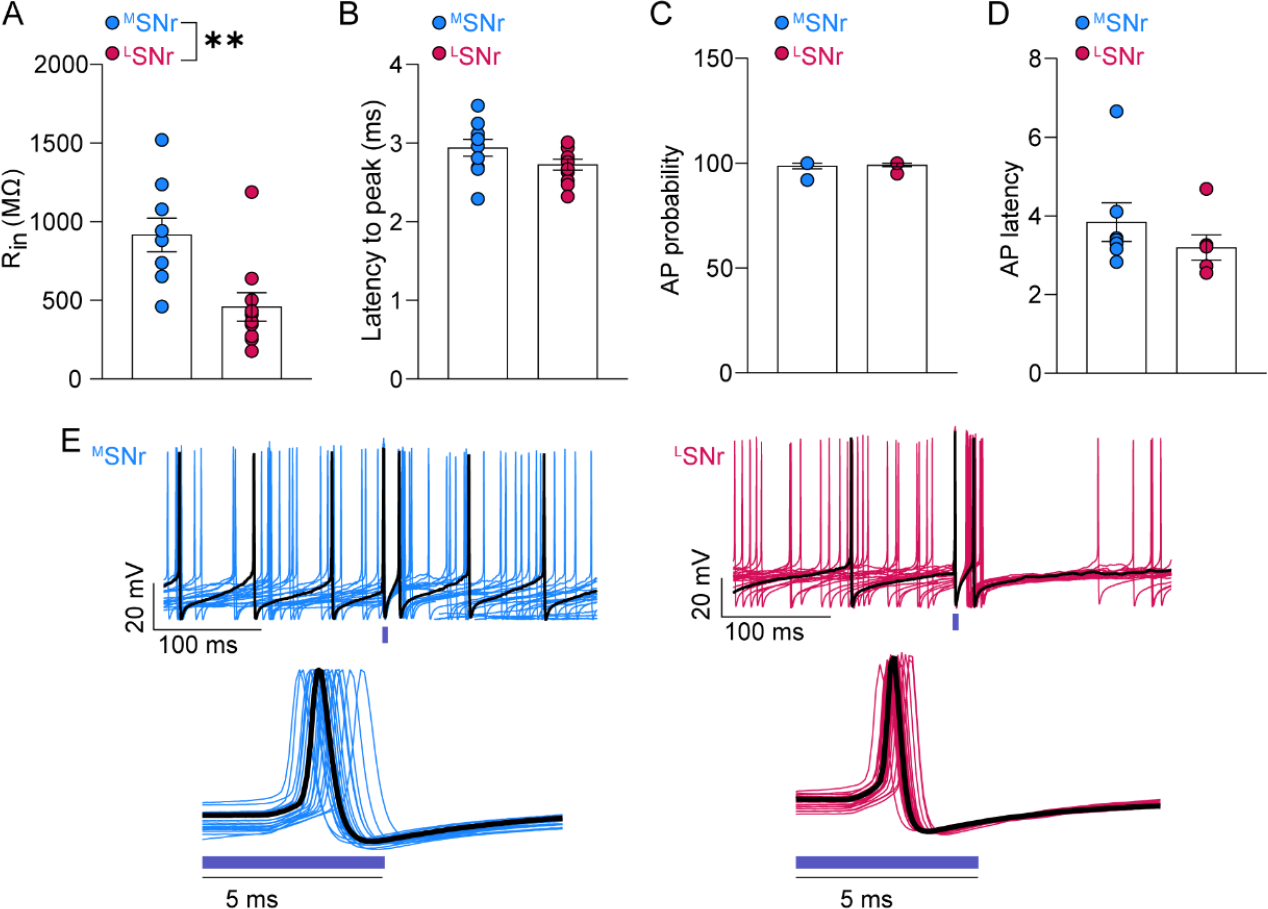
Functional and synaptic characterization of STN-SNr projections. **A**: Input resistance of ^M^SNr (N = 9 cells, 8 slices, 5 mice) and ^L^SNr (N = 10 cells, 10 slices, 6 mice) neurons recorded in fig. 2B-2C (Unpaired t-test, two-tailed, p = 0.0045). **B**: Latency to peak of oEPSCs recorded at STN-^M^SNr (N = 10 cells, 10 slices, 5 mice) and STN-^L^SNr (N = 11 cells, 10 slices, 6 mice) synapses. **C**: Probability of AP in ^M^SNr (N = 7 cells, 6 slices, 4 mice) and ^L^SNr (N = 6 cells, 6 slices, 3 mice) neurons upon STN input stimulation. **D**: Latency of AP in ^M^SNr (N = 7 cells, 6 slices, 4 mice) and ^L^SNr (N = 6 cells, 6 slices, 3 mice) neurons upon STN input stimulation. **E**: Example traces of ^M^SNr and ^L^SNr neuron firing during STN input stimulation.

## Notes

### Competing Interest Statement

The authors have declared no competing interest.

### Summary of Updates

The title and content of the manuscript have been updated

